# Shared and distinct genetic influences between cognitive domains and psychiatric disorder risk based on genome-wide data

**DOI:** 10.1101/2020.09.16.297408

**Authors:** CE Carey, Y Huang, RW Strong, 23andMe Research Team, S Aslibekyan, RC Gentleman, JW Smoller, JB Wilmer, EB Robinson, L Germine

## Abstract

Group-level cognitive performance differences are found in psychiatric disorders ranging from depression to autism to schizophrenia. To investigate the genetics of individual differences in fluid and crystallized cognitive abilities and their associations with psychiatric disorder risk, we conducted genome-wide association studies (GWAS) of a total of 335,227 consented 23andMe customers of European descent between the ages of 50 and 85, who completed at least one online test of crystallized cognitive ability (vocabulary knowledge, N=188,434) and/or fluid cognitive ability (visual change detection, N=158 888; digit-symbol substitution, N=132,807). All cognitive measures were significantly heritable (*h*^*2*^=0.10-0.16), and GWAS identified 25 novel genome-wide significant loci. Genetic correlation analyses highlight variable profiles of genetic relationships across tasks and disorders. While schizophrenia had moderate negative genetic correlations with tests of fluid cognition (visual change detection *r*_*g*_=−0.27, p<9.2e-24; digit-symbol substitution *r*_*g*_=−0.26, p<5.2e-27), it was only weakly negatively associated with crystalized cognition (vocabulary knowledge *r*_*g*_=−0.07, p<0.004). Autism, in contrast, showed a robust positive genetic correlation with vocabulary knowledge (*r*_*g*_=0.30, p<5.6e-13) and little to no genetic correlation with either fluid cognition task (*r*_*g*_’s<0.08, p’s>0.005). Crystalized and fluid cognitive abilities thus have correlated but distinct genetic architectures that relate to those of psychiatric disorders. Understanding the genetic underpinnings of specific cognitive abilities, and their relationships to psychiatric disorder risk, can inform the understanding of disease biology nosology and etiology.

## Introduction

Accumulating evidence indicates that psychiatric disorders may be conceptualized as maladaptive extremes of population distributions of information processing mechanisms (hereafter: cognition), rather than in terms of distinct diagnostic categories.^1–3^ Cognitive differences in psychiatric disorders are frequently detectable before disorder onset,^4,5^ found to a lesser degree in relatives of those with a diagnosis,^6,7^ and associated with symptom severity, recovery, and recurrence in prospective studies.^8^ Disruptions in specific cognitive processes are observed across disorders and linked to group-level differences in neural structure and function (e.g., cognition control^9^ and emotion regulation^10^), suggesting both shared and unique cross-disorder etiology.^11–13^ Variation in cognition thus provides a framework for understanding the biology, etiology, and nosology of psychiatric disorders.

Cognition has typically been divided into two broad components: fluid and crystallized abilities.^14–16^ Fluid cognitive abilities are associated with tasks that require speed or novel problem solving, particularly those where no previous knowledge is required (e.g., reaction time or working memory tests). In contrast, crystallized cognitive abilities are associated with tasks that require semantic knowledge accumulated through previous learning experiences (e.g., general information, arithmetic, or vocabulary tests). Although more nuanced theories of cognitive abilities have been proposed,^17,18^ the distinction between fluid and crystallized cognitive abilities is supported by factor analysis across a large literature that has identified distinct developmental trajectories,^19^ genetic architectures,^20^ and neural substrates.^21^

Understanding the genetic architecture of cognition and psychiatric disorders can provide clues about shared mechanisms that contribute to the etiology of these diseases.^22^ A substantial portion of risk for developing psychiatric disorders arises from genetic variation, with total heritability estimates as high as 80%,^23,24^ and common-variant heritability estimates of roughly 20%, for disorders such as schizophrenia and bipolar disorder.^25^ Genetic influences on psychiatric disorders, as with cognitive and neural disruptions, show substantial cross-disorder overlap, indicating that at least some genetic risk is not diagnosis-specific.^26–28^

Despite the explosion of psychiatric GWAS in recent years, the genetic architecture of specific aspects of cognition has remained largely unexplored. Though GWAS of individual cognitive mechanisms do exist (e.g., delay discounting,^29^ cognitive empathy,^30^ and executive function^31,32^), prior large-scale studies have focused primarily on a single measure of fluid cognitive abilities (e.g., in the UK Biobank^33^) or an aggregate “g” factor,^34–36^ or have used educational attainment (EA) as a proxy measure for cognition.^37–39^ The use of composite or proxy measures may obscure key differences in the genetic architecture of specific cognitive mechanisms and how they relate to disease. For example, the genetics of bipolar disorder and schizophrenia correlate positively with that of educational attainment, but negatively or nonsignificantly with that of aggregate cognitive measures;^40^ the use of EA as a proxy measure for cognition would thus lead to false conclusions about the disorder’s genetic relationship to cognition.

Prior investigations of polygenic risk for psychiatric disorders and associations with general vs. specific cognitive domains have in fact revealed disorder- and domain-specific associations not captured by general factors.^2,41–43^ Additionally, a prior study in the UK Biobank which assessed genetic associations between three individual measures of fluid cognition (i.e., verbal-numerical reasoning, reaction time, and memory)^44^ and indicators of physical and mental health revealed varying association profiles across neuropsychiatric disorders.^45^ However, whether genetic risk for neuropsychiatric disorders is differentially associated with fluid versus crystallized cognitive ability measures remains an open question, despite the large literature suggesting that these two aspects of cognition may have dissociable etiologies.

In the current study, we conducted genome-wide association analyses of a large population (N=335,227) of genotyped participants from the 23andMe personal genomics platform who participated in a suite of cognitive tests tapping fluid (i.e., processing speed and visual change detection) and crystallized (i.e., vocabulary knowledge) cognitive domains. We then assessed genetic overlap between six major neuropsychiatric disorders (i.e., schizophrenia, bipolar disorder, autism spectrum disorder, major depressive disorder, attention-deficit/hyperactivity disorder, and Alzheimer’s disease) and comparable cognitive and educational phenotypes (i.e., educational attainment and cognitive performance) with cognitive abilities in each domain in our study. These analyses provide insight into the genetic architecture of different cognitive processes and their relationships to each other and to psychiatric disease.

## Methods

### Participants

This study included 335,227 23andMe customers (58.6% female; *M*_*age*_=64.43[8.19]) who provided written informed consent and completed at least one of the three online cognitive assessments. Sampling was biased towards participants 50 and older as part of a larger 23andMe initiative. The study population was restricted to participants aged 50-85 at the time of assessment who were residing within the United States and determined to be of European ancestry based 23andMe’s proprietary algorithm.^46^ Due to the visual nature of the computer-based tasks, we excluded participants who self-reported serious and uncorrectable vision problems, including age-related macular degeneration, retinal vein occlusion, and retinitis pigmentosa. We also excluded participants who did not comply with the rules of each task, including those who walked away or skipped all trials. See **Table 1** for additional demographic information, including demographics split by cognitive assessment. The protocol governing human subjects research involving 23andMe participants was approved by the accredited Ethical and Independent Review Services (http://www.eandireview.com).

**Table 1.** Demographic information for individuals completing each of the 3 tasks.

|  | **FLICKER** | **VOCAB** | **DSST** |
| --- | --- | --- | --- |
| **N** | 158888 | 188434 | 132807 |
| **Age, M(SD)** | 64.2 (7.9) | 64.8 (8.4) | 64.2 (7.9) |
| **Female (%)** | 55.5% | 62.1% | 55.7% |
| **Years of Education M(SD)** | 16.2 (2.6) | 16.1 (2.6) | 16.1 (2.6) |
| **Missing, N(%)** | 5992 (3.7%) | 16796 (8.9%) | 5506 (4.1%) |
| **Type of Device** |  |  |  |
| **Laptops (%)** | 48.8% | NA | 48.6% |
| **Desktops (%)** | 51.2% | NA | 51.4% |

### Measures

#### Vocabulary Knowledge (Vocab)

Vocabulary Knowledge, or “Vocab,” is a test of verbal reasoning and long-term verbal memory, based on identification of word synonyms. In this test, which was modelled after the Wordsum task used in the General Social Surveys,^47^ participants have to select which of five words is most similar in meaning to a target word (**Supplementary Figure 1**). The primary outcome measure is the number of correct answers out of 20 questions. Internal split-half reliability in our sample was adequate (r=0.69). Vocabulary / synonym tests are widely used measures of crystallized cognitive ability, as they reflect accumulated word knowledge across the lifespan and are relatively insensitive to short-term changes in health.^48^ Similar to other tests of crystallized cognitive ability, performance tends to increase across early and middle adulthood with only modest decline in older age.^17^ Prior family studies have demonstrated moderate heritability (e.g., *h*^2^=50-63%) for measures of vocabulary knowledge.^49,50^

#### Flicker Change Detection (Flicker)

Flicker Change Detection, or “Flicker,” is a test of visual change detection, which loads on processing speed, visual search, and visual working memory aspects of fluid cognitive ability, and has been linked to neural activity in the frontoparietal networks.^51^ In this version, which is adapted from the Rensink change detection paradigm^52^ but with more tightly controlled stimuli, participants see a field of blue and yellow flashing dots (every 500ms; **Supplementary Figure 2**) where one dot is changing color. The participant presses the space bar as soon as they find the changing dot, and then indicate which dot was changing with their mouse. There are 2 training and 11 testing trials in total, and the primary outcome measure is median response time to accurately identify the changing dot on the 11 testing trials. Internal split-half reliability for this task in our sample was 0.49, which is low compared with traditional neuropsychological tasks, but similar to other brief tasks derived from the experimental literature for understanding individual differences.^53^ One prior preliminary study in twins using a similar task estimated heritability of task performance to be 52%.^54^

#### Digit-Symbol Substitution Test (DSST)

The Digit-Symbol Substitution Test, or “DSST,” was modeled on the digit symbol substitution test (WAIS IV^55^), a widely used measure of processing speed and fluid cognitive ability. It requires participants to match as many symbols and numbers as possible in 90 seconds based on a provided symbol/number key. In the current version, participants were provided with a key of 9 symbols, each matched with the numbers 1, 2, or 3 (where each number was matched with three symbols). The participant then responded using their keyboard to indicate which number matched the symbol shown in the center of their screen (**Supplementary Figure 3**). The primary outcome measure was the number of correct trials in 90 seconds. This is a validated measure of processing speed and short-term memory, with an internal split-half reliability in our sample of 0.92. Prior twin studies have demonstrated moderate heritability (e.g., *h*^2^=48-67%) for DSST performance.^56,57^

#### Genotyping, Imputation, and Quality Control

DNA extraction and genotyping of 23andMe participants were performed on saliva samples by the National Genetics Institute (NGI), a clinical laboratory improvement amendments (CLIA) licensed laboratory and a subsidiary of Laboratory Corporation of America. Imputation and quality control were conducted by 23andMe.

Saliva samples were genotyped on one of five Illumina genotyping platforms: HumanHap550+ Beadchip v1 and v2 platforms, OmniExpress+ BeadChip v3 platform, a customized array platform (v4), and the latest Illumina Infinium Global Screening array + customized array (v5) platform. Samples had minimum call rates of 98.5%. Individuals whose analyses failed repeatedly were re-contacted to provide additional samples. For extended genotyping and sample quality control details, see^58–60^.

Participant data for each genotyping platform was phased and imputed separately. As described previously,^29^ an internally-developed tool, Finch,^61^ was applied to generate phased participant data for the v1 to v4 platforms. For the X chromosome, separate haplotype graphs were built for the non-pseudoautosomal region and each pseudoautosomal region, which were phased separately. For the most recent v5 array, a similar approach was used with a new phasing algorithm, Eagle2.^62^ In preparation for imputation, first a single unified imputation reference panel was created by combining May 2015 release of the 1000 Genomes Phase 3 haplotypes^63^ with the UK10K imputation reference panel^64^ for better imputation performance in individuals of European descent.^65^ Second, each chromosome of the reference panel was split into segments of no more than 300,000 genotyped SNPs, with overlaps of 10,000 SNPs on each side. Finally, phased participant data was imputed against the merged reference panel using Minimac3,^66^ treating males as homozygous pseudodiploids for the non-pseudoautosomal region.

QC of genotyped and imputed SNPs was performed separately and then merged. SNPs genotyped on v1 and/or v2 platforms were excluded because of small sample size, as were those on the sex chromosomes due to unreliability. SNPs that failed a test for parent-offspring transmission were also removed using trio data. SNPs were further excluded with a Hardy-Weinberg p<10^−20^, a call rate of <90%, substantial sex effect (r^2^>0.1 by ANOVA), significant genotyping date effects (p<10^−50^ by ANOVA), or probes matching multiple genomic positions in the reference genome (‘self chain’). Imputed SNPs were excluded for having an average imputation *r*^2^<0.3 or a strong batch effect (p<10^−50^ by ANOVA). Across all SNPs, we additionally excluded SNPs with an available sample size of less than 20% of the total sample size as well as those with MAF<0.1%. Post-QC, associations tests were performed on 13,806,893 SNPs for Vocab and on 13,806,898 SNPs for Flicker and DSST.

### Statistical Analyses

#### Association Analyses

Using the 23andMe internal pipeline,^29^ three genome-wide association studies (GWAS) were conducted among participants of European descent. For each, a maximal set of unrelated individuals was chosen using a segmental identity-by-descent (IBD) estimation algorithm.^67^

Association tests were performed using linear regression and assuming additive allelic effects. Outcomes were defined as: 1) number of incorrect answers out of 20 questions for Vocab (square root transformed), 2) median response time of correct responses for Flicker (log-transformed), and 3) number of correct trials in 90 seconds for DSST. When reporting all association results, the sign of regression coefficients was flipped for Vocab and Flicker such that a positive effect reflects better performance (i.e., less incorrect answers for Vocab and quicker response time for Flicker). Covariates included age (years), sex (male vs. female), age*sex interaction, 10 genetic principal components (PC), and genotyping platform. For both Flicker and DSST, we additionally adjusted for device (desktop vs. laptop) due to its significant influence on performance.

#### Heritability and Genetic Correlation

SNP-based heritability for each measure was calculated using LD score regression^68^ and the resultant summary statistics from each GWAS. Genetic correlations among these measures and with additional phenotypes (i.e., psychiatric disorders,^25,69–72^ Alzheimer's disease,^73^ educational attainment,^38^ and cognitive performance;^38^ **Supplementary Table 1**) were estimated using LDSR as well.

## Results

### Phenotypic Assessment

A total of 335,227 eligible participants met inclusion criteria: 188,434 for Vocab, 158,888 for Flicker, and 132,807 for DSST. The average number of correct responses on Vocab was 16.7(2.5); average median response time to correctly identify the changing dot on Flicker across individuals (in milliseconds) was 8884(5136); and the average number of correct trials within 90 seconds on DSST was 49.6(10.8). Among individuals who completed more than one cognitive task, performance was correlated across tasks when adjusted for all GWAS covariates, with both fluid cognitive ability measures correlated more strongly with each other (r_Flicker-DSST_=0.33, p<1e-300, N=91,774) than with the single measure of crystallized cognitive ability (r_Flicker-Vocab_ =0.10, p=9.48e-102, N=47,321; r_DSST-Vocab_=0.16, p=1.45e-217, N=35,976). Performance on all three measures was consistent with previous publications^74^ and data from Testmybrain.org,^53^ with similar associations between performance and demographic characteristics.

In the context of the null GWAS model, increased age was associated with better performance on Vocab (β=0.007, p=1.7e-96), but worse performance on Flicker (β=−0.02, p<10^−300^) and DSST (β=−0.72, p<10^−300^). Female sex was associated with poorer performance on all tasks (β_Vocab_=−0.39, p=7.4e-40; β_Flicker_=—0.18, p=4.1e-21; β_DSST_=−2.88, p=7.0e-12). Use of a laptop vs. desktop computer was associated with better performance on DSST (β=0.36, p=7.6e-13) but worse performance on Flicker (β=−0.05, p=5.9e-94). Full association results for each null GWAS model are reported in **Supplementary Tables 2-4**.

### Heritability

All three tests demonstrated significant SNP-based heritability as estimated using LDSR (Vocab: *h^2^*=0.101[0.005], Flicker: *h*^2^=0.120[0.006], DSST: *h^2^*=0.161[0.007]; all z>20).

### Genome-wide Association Results

Genome-wide association analyses yielded 11 novel significant (i.e., p<5e-8) loci for Vocab (λ= 1.199, λ_2000_=1.002), and 7 loci each for Flicker (λ= 1.196, λ_2000_=1.002) and DSST (λ= 1.219, λ_2000_=1.003; **Table 2**; **Figure 1**; see **Supplementary Tables 5-7**for all loci p<1e-6; see **Supplementary Figures 4-6** for QQ plots).

**Table 2.**
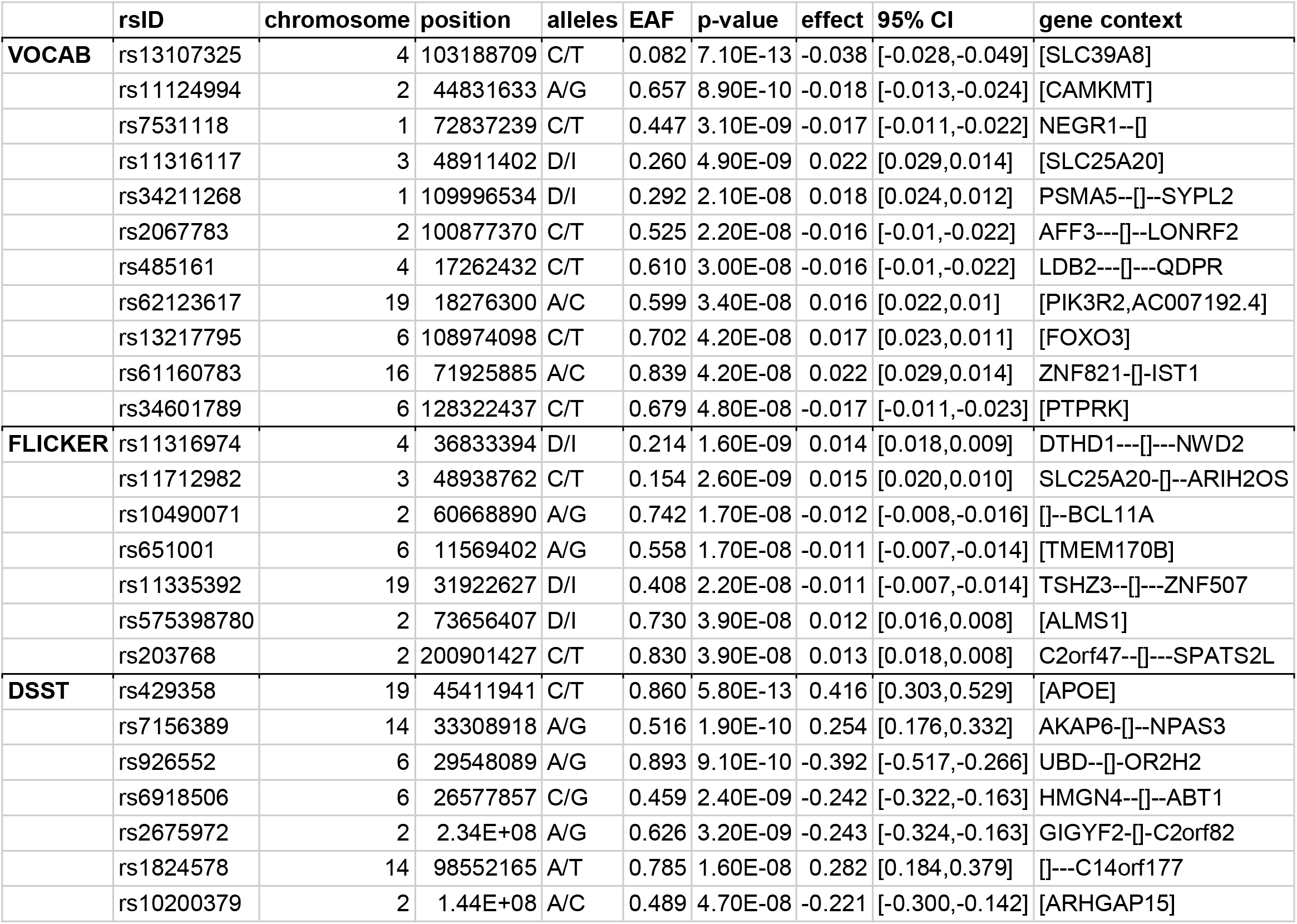
Genome-wide significant loci for each task 26

**Figure 1.**
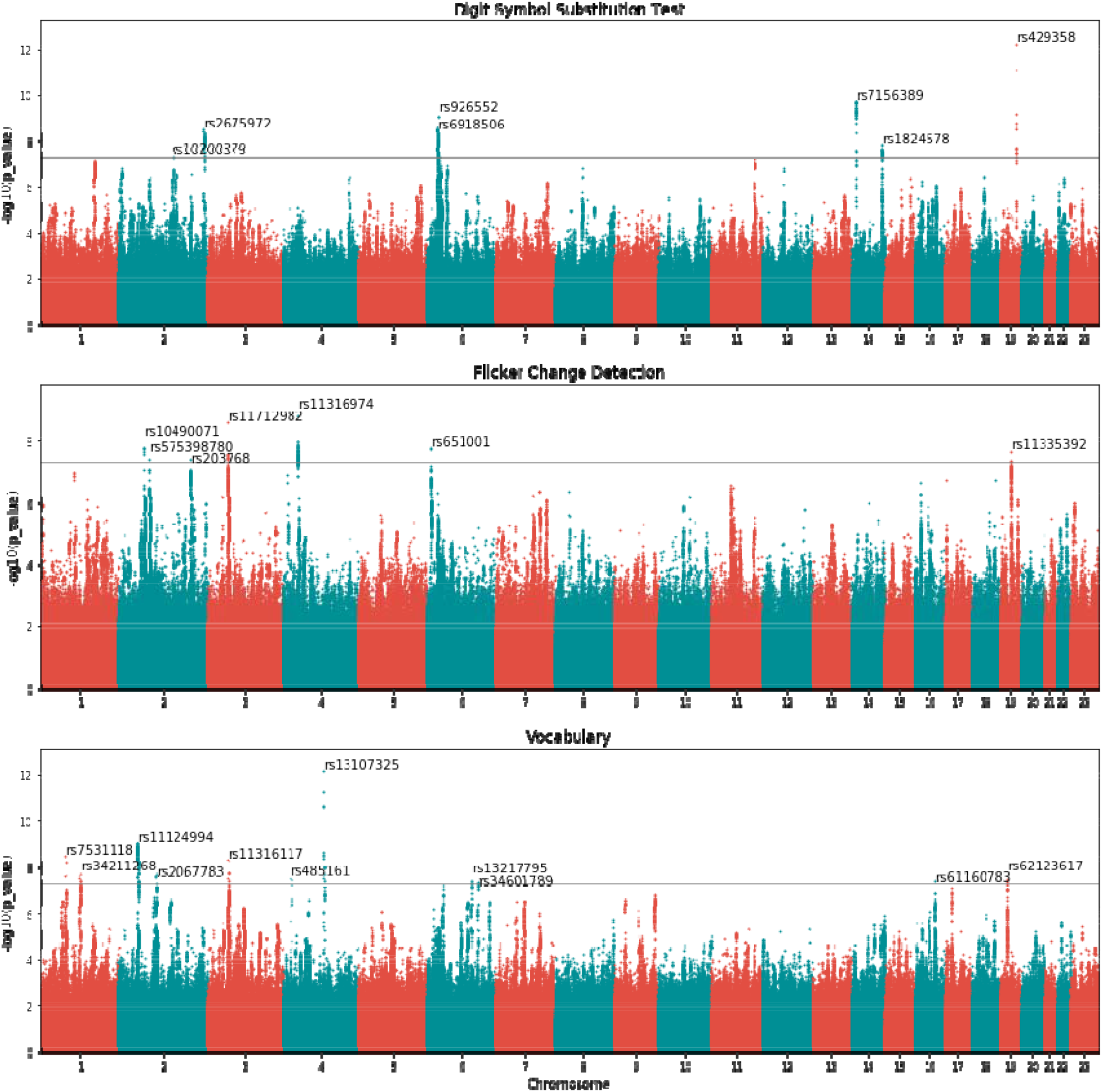
Manhattan plots for all GWAS performed in this study

The top hit for Vocab, rs13107325 (p=7.1e-13; **Supplementary Figure 7**), is a missense variant within *SLC39A8* that alters the amino acid at position 391. The top SNP for Flicker, rs11316974 (p=1.6e-09; **Supplementary Figure 8**), is an indel in an intergenic region between the genes *DTHD1* and *NWD2* that has not been previously associated with any traits. The top SNP for DSST, rs429358 (p=5.80E-13; **Supplementary Figure 9**), is a missense variant within *APOE* that alters the amino acid at position 130 and, in doing so, greatly increases the risk of Alzheimer’s disease.^75^

### Genetic Correlation

Genetic correlation among the three cognition measures was moderate (*r*_*g*_=0.247-0.650), with a noticeably higher correlation between the two fluid cognitive ability measures (*r*_*g*_=0.650) than with the single measure of crystallized cognitive ability (*r*_*g_Flicker*_=0.247, *r*_*g_DSST*_=0.308). Using out-of-sample EA and CP summary statistics, fluid ability measures were more highly related to CP (*r*_*g_Flicker*_=0.514, *r_g_DSST_*=0.562) than to EA (*r*_*g_Flicker*_=0.163, *r*_*g_DSST*_=0.317); vocabulary knowledge genetically correlated to both EA and CP at a similar magnitude (*r*_*g_EA*_=0.659, *r*_*g_CP*_=0.724).

Genetic correlations with neuropsychiatric disorders revealed generally widespread negative associations with cognitive performance across domains, with some notable exceptions (**Table 3**). DSST performance was negatively associated with genetic risk for each neuropsychiatric disorder tested (*r*_*g*_=−0.187 to −0.265), with the exception of ASD (*r*_*g*_=0.024). Performance on Flicker, in contrast, was negatively genetically correlated with only BIP (*r*_*g*_=−0.183) and SCZ (*r*_*g*_=−0.269). Vocabulary knowledge was negatively associated with ADHD risk (*r*_*g*_=−0.234), but positively associated with ASD risk (*r*_*g*_=0.301).

**Table 3.** Genetic correlations with published summary statistics

|  | Vocab |  | Flicker |  | DSST |  |
| --- | --- | --- | --- | --- | --- | --- |
| trait/disorder | rg | p-value | rg | p-value | rg | p-value |
| attention deficit hyperactivity disorder | **-0.234** | 1.19E-11 | -0.047 | 1.94E-01 | **-0.208** | 3.23E-09 |
| autism spectrum disorders | **0.301** | 7.37E-13 | 0.079 | 4.37E-02 | 0.024 | 5.48E-01 |
| bipolar disorder | 0.048 | 1.90E-01 | **-0.183** | 2.90E-08 | **-0.187** | 1.24E-09 |
| major depressive disorder | -0.071 | 5.06E-02 | -0.105 | 2.88E-03 | **-0.222** | 2.72E-12 |
| schizophrenia | -0.074 | 2.63E-03 | **-0.269** | 9.19E-24 | **-0.265** | 1.02E-26 |
| Alzheimer's disease | -0.111 | 5.63E-02 | -0.041 | 4.45E-01 | **-0.221** | 9.90E-05 |
| educational attainment | **0.659** | 2.87E-278 | **0.163** | 1.58E-14 | **0.317** | 7.36E-65 |
| cognitive performance | **0.724** | 0.00E+00 | **0.514** | 2.83E-118 | **0.562** | 2.83E-170 |

Bold values indicate significant correlations after correction for multiple testing

## Discussion

Here, we report results from the largest study to date on the genetics of fluid and crystallized cognitive abilities with respect to genetic risk for neuropsychiatric disorders. We conducted GWAS of performance on three cognitive tests tapping crystallized (i.e., vocabulary knowledge) and fluid (i.e., processing speed, visual change detection) cognitive abilities, based on data from 335,227 genotyped 23andMe participants. These analyses 1) demonstrate heritability for and patterns of genetic overlap between all cognitive domains tested; 2) identify novel, biologically informative loci for each; and 3) reveal unique profiles of genetic association with neuropsychiatric disorders.

Heritability analyses of all three measures confirm that they are influenced by common variants (*h*^2^=0.10-0.16) to a similar degree as other complex traits such as educational attainment (EA; *h*^2^=0.147^38^) and neuroticism (*h^2^*=0.100^76^). Furthermore, patterns of genetic overlap across measures are consistent with classical theories of intelligence proposing both a central “g” factor as well as separate components for fluid and crystallized abilities.^14–16^ Notably, the two fluid cognition measures (i.e., flicker change detection [Flicker] and the digit-symbol substitution test [DSST]) were more genetically correlated with each other (*r*_*g*_=0.650) than with the single measure of crystallized cognitive ability (i.e., vocabulary knowledge [Vocab]; *r_g_Flicker_*=0.247, *r_g_DSST_*=0.308). This pattern was further supported by genetic associations with EA and cognitive performance (CP),^38^ in which the two fluid measures were more highly related to CP (*r_g_Flicker_*=0.514, *r_g_DSST_*=0.562) than to EA (*r_g_Flicker_*=0.163, *r_g_DSST_*=0.317). Overall, while genetic overlap between the three measures (minimum *r*_*g*_=0.247) supports the theory of a shared underlying etiology or “g” factor, incomplete genetic overlap between measures (maximum *r*_*g*_=0.650), along with different magnitudes of genetic correlation between fluid and crystallized measures, suggests that existing GWAS approaches that combine across measures and/or use EA as a proxy for cognitive abilities might obscure the unique genetic contributions to different facets of cognition.

The consequences of this incomplete pattern of genetic overlap can be seen in an examination of the top loci for each measure in our study, which revealed no common significant or suggestive loci across measures. The top hit for Vocab, rs13107325 (p=7.1e-13), is a missense variant within *SLC39A8* that alters the amino acid at position 391 that has been previously linked to a host of complex traits including intelligence,^36^ autism,^77^ and schizophrenia^69,78^ and has been functionally associated with altered glycosylation in the brain, which is critical to neurodevelopment.^79^ Similarly, the top hit for DSST, rs429358 (p=5.80E-13), is a known pathogenic missense variant within *APOE* that alters the amino acid at position 130 and, in doing so, greatly increases the risk of Alzheimer’s disease.^75^ It and nearby variants in high linkage disequilibrium (LD; r^2^>0.70) have also been linked to biomarkers of Alzheimer’s disease including tau^80^ and amyloid-β levels.^81^ In addition to Alzheimer’s disease, it has been previously associated with measures of age-related cognitive decline^82^ and verbal memory.^83,84^ The top SNP for Flicker, rs11316974 (p=1.6e-09), is an indel in an intergenic region between the genes *DTHD1* and *NWD2* that has not been previously associated with any traits. Among the other significant variants across the three measures, many were in high LD with variants previously linked to relevant phenotypes such as general cognitive function,^39,8537^ intelligence,^36^ educational attainment,^86^ schizophrenia,^69^ and autism.^77^ The lack of overlap in top loci across domains, combined with the known neural and prior genetic associations of these loci, suggest that parsing the genetic influences on different domains of cognition could prove fruitful in providing mechanistic insight.

Furthermore, aside from examining the top loci, genetic correlation analyses with GWAS of six major neuropsychiatric disorders^25,69–73^ reveal robust, dissociable association patterns for autism versus bipolar disorder and schizophrenia, with 1) moderate positive genetic correlations between autism and crystallized (*r*_*g*_=0.301), but not fluid (*r*_*g*_=0.024 to 0.080, all p’s>0.04), cognitive abilities; and 2) significant negative genetic correlations between both schizophrenia and bipolar disorder and fluid (*r*_*g*_=−0.187 to −0.269), but not crystallized (*r*_*g*_=−0.074 to 0.048, all p’s>0.002), abilities. These differential associations are notable given consistent positive genetic associations,^40,45^ but negative diagnostic associations (perhaps with the exception of BIP^87^),^88,89^ between these disorders and EA. Our results suggest that the etiology of schizophrenia and bipolar disorder involves the impairment of fluid cognitive abilities, while genetically conferred risk for autism generally spares fluid cognitive abilities and in fact confers greater potential to accumulate knowledge (here: verbal knowledge). These findings are consistent with prior studies suggesting that the genetics underlying autism and schizophrenia have distinct influences on neural and cognitive development.^43,90^ Understanding the genetic architecture of fluid and crystallized abilities, and pathways that differentially contribute to each, thus has the potential to provide greater clarity on the mechanisms that influence development of these disorders.

## Limitations

A noteworthy limitation of the current study is sample ascertainment and inclusion criteria. Our age range was restricted (i.e., 50-85), and, as such, the results may not generalize to other age strata or may be contaminated by cognitive decline. This is particularly true of the fluid cognition measures, performance on which is known to decline with age,^17,91^ versus the crystallized cognition measure, which has been shown to be relatively immune to the effects of aging.^17^ Notably, the top locus for our measure of processing speed is a known marker for Alzheimer’s disease that has been previously linked to age-related cognitive decline.^82^ However, the use of a restricted age range is not unique to our study; notably, the age range of the UK Biobank, in which previous GWAS of specific cognitive abilities have been conducted,^31,44,45^ is similarly restricted to 40-70 at recruitment.^33^ Furthermore, while the 23andMe platform provides an engaging means of participant recruitment for genetics research, participants self-selection may bias results. For example, older adults with lower cognitive functioning are unlikely to have participated in the study due to the reliance on web-based recruitment, engagement, and testing. Similarly, participants from lower socioeconomic status groups are unlikely to have paid to receive personalized genetic results from 23andMe. While the impact of such biases on our results is difficult to quantify, it is reassuring that our results are consistent with a prior study of genetic correlations between cognitive measures and neuropsychiatric disorders in the UK Biobank,^45^ and also that we observed expected genetic correlations with cognitive performance and educational attainment as estimated in other, younger and non-age-restricted samples.^38^

## Conclusions & Future Directions

Our results build on a growing literature investigating the genetic architecture of cognitive abilities and their relationship to neuropsychiatric disorders. While we propose and provide evidence for a general distinction between fluid and crystallized cognition, our measures also differ in their reliance on visual versus verbal information, and measurement of speed versus accuracy as the primary outcome of interest. It is possible that these distinctions are meaningfully associated with disease mechanisms; future work extending these analyses to other objectively measured cognitive domains would clarify which are most informative. Future studies might also investigate specific variants and pathways driving the genetic overlap across measures and disorders, as well as how they relate to neural structure and function. Overall, our results argue for a more nuanced treatment of cognitive abilities in psychiatric genetics that reflects the architecture of cognition beyond the general “g” factor. In an era where digital cognitive assessment is low-cost and accessible, achieving the sample sizes necessary to generate new knowledge at the intersection of psychiatric and cognitive genetics is important and feasible.

## Acknowledgements

CEC is funded by 5R01MH111813 (PI Robinson). This work was funded by a Milken Institute grant to 23andMe, Inc.

We would also like to thank the following members of the 23andMe Research Team: Michelle Agee, Babak Alipanahi, Adam Auton, Robert K. Bell, Katarzyna Bryc, Dennis Byrne, Sarah L. Elson, Anna Faaborg, Pierre Fontanillas, Nicholas A. Furlotte, Pooja Gandhi, Travis Hairfield, Eric Hall, Barry Hicks, David A. Hinds, Karen E. Huber, Ethan M. Jewett, Yunxuan Jiang, Aaron Kleinman, Keng-Han Lin, Nadia K. Litterman, Bo Lopker, Jennifer C. McCreight, Matthew H. McIntyre, Kimberly F. McManus, Joanna L. Mountain, Elizabeth S. Noblin, Carrie A.M. Northover, Steven J. Pitts, G. David Poznik, J. Fah Sathirapongsasuti, Janie F. Shelton, Suyash Shringarpure, Chao Tian, Joyce Y. Tung, Vladimir Vacic, Xin Wang, Catherine Weldon, and Catherine H. Wilson.

