## Supplemental Materials for "Shared and distinct genetic influences between cognitive domains and psychiatric disorder risk based on genome-wide data"

^3^23andMe, Sunnyvale, CA, USA

^4^Institute for Technology in Psychiatry, McLean Hospital, Belmont, MA, USA

^5^Department of Psychiatry, Harvard Medical School, Boston, MA, USA

^6^Psychiatric and Neurodevelopmental Unit, Massachusetts General Hospital, Boston, MA, USA

^7^Department of Psychology, Wellesley College, Wellesley, MA, USA

^8^Department of Epidemiology, Harvard T.H. Chan School of Public Health, Boston, MA, USA

**Corresponding Author:**

Laura Germine

Institute for Technology in Psychiatry
McLean Hospital

115 Mill Street

Oaks 358A
Belmont, MA, 02478

[Supplementary Table 5. Non-significant suggestive loci (p<1e-6) for Vocabulary Knowledge	7](#_Toc51026004)

[Supplementary Table 6. Non-significant suggestive loci (p<1e-6) for Flicker Change Detection	8](#_Toc51026005)

[Supplementary Table 7. Non-significant suggestive loci (p<1e-6) for the Digit Symbol Substitution Test	9](#_Toc51026006)

### Supplementary Tables

#### Supplementary Table 1. Published summary statistics used for genetic correlation analyses

| **trait/disorder** | **N** | **N_case** | **N_control** | **h2_SNP** | **citation** |
| --- | --- | --- | --- | --- | --- |
| attention deficit hyperactivity disorder | 53,293 | 19,099 | 34,194 | 0.216 | Demontis et al., 2019 |
| autism spectrum disorders | 46,351 | 18,382 | 27,969 | 0.118 | Grove et al., 2019 |
| bipolar disorder | 41,653 | 20,129 | 21,524 | 0.195 | Ruderfer et al., 2018 |
| major depressive disorder | 173,005 | 59,851 | 113,154 | 0.089 | Wray et al., 2018 |
| Schizophrenia | 77,096 | 33,640 | 43,456 | 0.249 | Ripke et al., 2014 |
| Alzheimer's disease | 455,258 | 71,880 | 383,378 | 0.055 | Jansen et al., 2019 |
| educational attainment | 1,131,881 |  |  | 0.147 | Lee et al., 2018 |
| cognitive performance | 257,841 |  |  | 0.199 | Lee et al., 2018 |

#### Supplementary Table 2. Null GWAS model association results for Vocabulary Knowledge

| **Covariate** | **Estimate** | **Std. Error** | **t value** | **Pr(>\|t\|)** |
| --- | --- | --- | --- | --- |
| Age | 0.007 | 0.000 | 21 | 1.70E-96 |
| sex: female | -0.386 | 0.029 | -13 | 7.40E-40 |
| PC1 | -0.024 | 0.002 | -13 | 5.20E-40 |
| PC2 | 0.007 | 0.002 | 4 | 2.70E-04 |
| PC3 | 0.045 | 0.002 | 24 | 3.80E-130 |
| PC4 | 0.009 | 0.002 | 5 | 1.60E-06 |
| PC5 | -0.002 | 0.002 | -1 | 2.20E-01 |
| PC6 | -0.002 | 0.002 | -1 | 3.70E-01 |
| PC7 | 0.006 | 0.002 | 3.2 | 1.30E-03 |
| PC8 | -0.001 | 0.002 | -0.6 | 5.20E-01 |
| PC9 | 0.004 | 0.002 | 2.2 | 2.50E-02 |
| PC10 | -0.002 | 0.002 | -1 | 3.00E-01 |
| v3_1_platformv3.1 | -0.043 | 0.016 | -2.6 | 8.50E-03 |
| v4_platformv4 | -0.172 | 0.013 | -13.1 | 3.00E-39 |
| v5_platformv5 | -0.337 | 0.013 | -25.6 | 9.30E-145 |
| age*sex | 0.006 | 0.000 | 14.1 | 6.70E-45 |

#### Supplementary Table 3. Null GWAS model association results for Flicker Change Detection

| **Covariate** | **Estimate** | **Std. Error** | **t value** | **Pr(>\|t\|)** |
| --- | --- | --- | --- | --- |
| Age | -0.025 | 0.000 | -115 | 0.00E+00 |
| sex: female | -0.179 | 0.019 | -9 | 4.10E-21 |
| PC1 | 0.004 | 0.001 | 4 | 2.50E-04 |
| PC2 | 0.001 | 0.001 | 1 | 2.20E-01 |
| PC3 | 0.004 | 0.001 | 4 | 5.60E-04 |
| PC4 | -0.015 | 0.001 | -13 | 4.60E-36 |
| PC5 | -0.011 | 0.001 | -10 | 2.50E-21 |
| PC6 | -0.002 | 0.001 | -2 | 1.10E-01 |
| PC7 | 0.000 | 0.001 | 0.1 | 9.10E-01 |
| PC8 | 0.003 | 0.001 | 2.3 | 2.10E-02 |
| PC9 | -0.001 | 0.001 | -0.8 | 4.40E-01 |
| PC10 | 0.002 | 0.001 | 2 | 4.60E-02 |
| v3_1_platformv3.1 | -0.014 | 0.013 | -1.1 | 2.70E-01 |
| v4_platformv4 | -0.048 | 0.010 | -4.6 | 3.60E-06 |
| v5_platformv5 | -0.100 | 0.010 | -9.7 | 2.10E-22 |
| age*sex | 0.003 | 0.000 | 8.7 | 4.70E-18 |
| device: laptop | -0.048 | 0.002 | -20.6 | 5.90E-94 |

#### Supplementary Table 4. Null GWAS model association results for the Digit Symbol Substitution Test

| **Covariate** | **Estimate** | **Std. Error** | **t value** | **Pr(>\|t\|)** |
| --- | --- | --- | --- | --- |
| age | -0.717 | 0.005 | -150 | 0.00E+00 |
| sex: female | -2.882 | 0.420 | -7 | 7.00E-12 |
| PC1 | -0.192 | 0.026 | -8 | 6.00E-14 |
| PC2 | -0.117 | 0.026 | -5 | 5.50E-06 |
| PC3 | 0.231 | 0.026 | 9 | 1.80E-19 |
| PC4 | -0.197 | 0.026 | -8 | 1.60E-14 |
| PC5 | -0.163 | 0.026 | -6 | 3.80E-10 |
| PC6 | -0.042 | 0.025 | -2 | 9.90E-02 |
| PC7 | 0.023 | 0.025 | 0.9 | 3.60E-01 |
| PC8 | -0.022 | 0.025 | -0.9 | 3.80E-01 |
| PC9 | 0.004 | 0.025 | 0.2 | 8.80E-01 |
| PC10 | 0.044 | 0.025 | 1.7 | 8.10E-02 |
| v3_1_platformv3.1 | -0.594 | 0.300 | -2 | 4.80E-02 |
| v4_platformv4 | -1.589 | 0.245 | -6.5 | 8.80E-11 |
| v5_platformv5 | -1.933 | 0.241 | -8 | 1.10E-15 |
| age*sex | 0.087 | 0.006 | 13.3 | 1.50E-40 |
| device: laptop | 0.367 | 0.051 | 7.2 | 7.60E-13 |

#### Supplementary Table 5. Non-significant suggestive loci (p<1e-6) for Vocabulary Knowledge

| **rsID** | **chromosome** | **position** | **alleles** | **EAF** | **p-value** | **effect** | **95% CI** | **gene context** |
| --- | --- | --- | --- | --- | --- | --- | --- | --- |
| rs149708654 | 6 | 33260187 | D/I | 0.832 | 6.90E-08 | 0.021 | [0.028,0.013] | [RGL2] |
| rs8069321 | 17 | 18117610 | C/T | 0.380 | 9.10E-08 | -0.016 | [-0.01,-0.021] | ALKBH5-[]--LLGL1 |
| rs1281159 | 9 | 129858518 | C/T | 0.475 | 1.70E-07 | -0.015 | [-0.009,-0.021] | [RALGPS1,ANGPTL2] |
| rs33950927 | 2 | 82062113 | D/I | 0.692 | 2.20E-07 | 0.016 | [0.022,0.01] | [] |
| esv3620013 | 9 | 23370987 | X/Y | 0.409 | 2.60E-07 | 0.015 | [0.021,0.009] | DMRTA1---[]---ELAVL2 |
| rs5834597 | 2 | 138742773 | D/I | 0.782 | 2.70E-07 | -0.018 | [-0.011,-0.025] | [HNMT] |
| rs13113989 | 4 | 65764491 | C/G | 0.537 | 2.80E-07 | -0.015 | [-0.009,-0.021] | TECRL---[]---EPHA5 |
| rs1699445 | 7 | 68778574 | C/T | 0.334 | 3.00E-07 | -0.016 | [-0.01,-0.022] | []---AUTS2 |
| rs11207639 | 1 | 61051901 | C/T | 0.358 | 3.00E-07 | 0.015 | [0.021,0.009] | C1orf87---[]---NFIA |
| rs12205803 | 6 | 157215580 | A/G | 0.250 | 3.40E-07 | -0.017 | [-0.01,-0.023] | [ARID1B] |
| rs3735478 | 7 | 44800176 | G/T | 0.287 | 3.40E-07 | 0.017 | [0.024,0.011] | [ZMIZ2] |
| rs3004179 | 6 | 98513032 | A/G | 0.597 | 3.60E-07 | -0.015 | [-0.009,-0.021] | MMS22L---[]---POU3F2 |
| rs62254271 | 3 | 53170191 | C/T | 0.049 | 4.80E-07 | -0.035 | [-0.021,-0.048] | RFT1-[]--PRKCD |
| rs35405441 | 3 | 94850828 | D/I | 0.582 | 5.40E-07 | -0.014 | [-0.009,-0.02] | [] |
| rs9383256 | 6 | 17170871 | C/T | 0.703 | 6.80E-07 | 0.016 | [0.022,0.01] | STMND1--[]---RBM24 |
| rs2814726 | 9 | 88014646 | C/T | 0.238 | 6.90E-07 | 0.017 | [0.023,0.01] | NTRK2---[]---AGTPBP1 |
| rs146474368 | 16 | 56312736 | C/T | 0.001 | 7.90E-07 | 0.262 | [0.365,0.158] | [GNAO1] |
| rs144212148 | 5 | 62585434 | A/G | 0.004 | 8.10E-07 | 0.147 | [0.205,0.088] | IPO11---[]---HTR1A |
| rs9640774 | 7 | 116447608 | C/T | 0.588 | 9.60E-07 | 0.014 | [0.02,0.009] | MET--[]--CAPZA2 |

#### Supplementary Table 6. Non-significant suggestive loci (p<1e-6) for Flicker Change Detection

| **rsID** | **chromosome** | **position** | **alleles** | **EAF** | **p-value** | **effect** | **95% CI** | **gene context** |
| --- | --- | --- | --- | --- | --- | --- | --- | --- |
| rs2678897 | 2 | 58169418 | A/G | 0.390 | 9.40E-08 | 0.01 | [0.014,0.006] | [VRK2] |
| rs6682836 | 1 | 97591557 | C/G | 0.528 | 1.10E-07 | 0.01 | [0.013,0.006] | [DPYD] |
| rs13110734:G | 4 | 10727529 | C/G | 0.419 | 1.40E-07 | 0.01 | [0.014,0.006] | CLNK--[]---HS3ST1 |
| rs4790293 | 17 | 1841742 | C/T | 0.633 | 1.90E-07 | -0.01 | [-0.006,-0.014] | [RTN4RL1] |
| rs199904655 | 18 | 67324700 | D/I | 0.976 | 2.00E-07 | 0.032 | [0.043,0.020] | [DOK6] |
| rs113409737 | 16 | 10137830 | D/I | 0.691 | 2.40E-07 | 0.01 | [0.014,0.006] | [GRIN2A] |
| rs397848056 | 11 | 46838338 | D/I | 0.124 | 2.80E-07 | 0.014 | [0.020,0.009] | [CKAP5] |
| rs549033344 | 11 | 57343227 | D/I | 0.818 | 3.50E-07 | -0.012 | [-0.007,-0.017] | UBE2L6-[]--SERPING1 |
| rs60345653 | 8 | 22101716 | A/G | 0.312 | 4.50E-07 | 0.01 | [0.014,0.006] | PHYHIP--[]-POLR3D |
| rs62474715 | 7 | 115061169 | G/T | 0.188 | 4.60E-07 | 0.012 | [0.016,0.007] | MDFIC---[]---TFEC |
| rs112710530 | 6 | 26700034 | D/I | 0.535 | 5.10E-07 | -0.011 | [-0.007,-0.015] | ZNF322--[]---HIST1H2BJ |
| rs372300621 | 7 | 97948860 | D/I | 0.413 | 6.30E-07 | -0.01 | [-0.006,-0.014] | [BAIAP2L1] |
| rs6786794 | 3 | 31533201 | C/G | 0.456 | 6.60E-07 | 0.009 | [0.013,0.006] | GADL1---[]--STT3B |
| rs12217221 | 10 | 79396463 | C/T | 0.238 | 7.00E-07 | 0.011 | [0.015,0.007] | [KCNMA1] |
| rs113205442 | 19 | 50085427 | A/G | 0.813 | 7.80E-07 | -0.012 | [-0.007,-0.017] | [PRRG2] |
| rs6467499 | 7 | 133348922 | G/T | 0.538 | 8.20E-07 | -0.009 | [-0.005,-0.012] | [EXOC4] |
| rs537064043 | 1 | 155212410 | D/I | 0.253 | 8.30E-07 | -0.011 | [-0.006,-0.015] | [GBA] |

#### Supplementary Table 7. Non-significant suggestive loci (p<1e-6) for the Digit Symbol Substitution Test

| **rsID** | **chromosome** | **position** | **alleles** | **EAF** | **p-value** | **effect** | **95% CI** | **gene context** |
| --- | --- | --- | --- | --- | --- | --- | --- | --- |
| rs11214609 | 11 | 113316102 | C/G | 0.392 | 6.20E-08 | -0.226 | [-0.308,-0.144] | [DRD2] |
| rs6700292 | 1 | 176134531 | G/T | 0.593 | 7.70E-08 | -0.234 | [-0.319,-0.149] | [RFWD2] |
| rs78903320 | 6 | 43179655 | C/T | 0.911 | 1.30E-07 | -0.368 | [-0.505,-0.232] | [CUL9] |
| rs6982915 | 8 | 64842662 | A/G | 0.242 | 1.50E-07 | 0.244 | [0.153,0.335] | YTHDF3---[]---BHLHE22 |
| rs61937595 | 12 | 57682956 | C/T | 0.084 | 1.60E-07 | 0.404 | [0.253,0.555] | [RP11-123K3.4,R3HDM2] |
| rs78959944 | 2 | 4874702 | A/C | 0.789 | 1.60E-07 | -0.262 | [-0.361,-0.164] | DCDC2C---[]---SOX11 |
| rs4583406 | 2 | 147730154 | C/T | 0.253 | 2.00E-07 | -0.243 | [-0.334,-0.151] | []---ACVR2A |
| rs551444803 | 2 | 201284434 | D/I | 0.783 | 2.90E-07 | 0.268 | [0.166,0.371] | [SPATS2L] |
| rs2937607 | 2 | 155727726 | C/T | 0.735 | 3.00E-07 | 0.233 | [0.144,0.323] | KCNJ3--[] |
| rs4800012 | 18 | 35798338 | A/C | 0.481 | 3.70E-07 | 0.203 | [0.125,0.281] | CELF4---[] |
| rs6553441 | 4 | 170448464 | C/T | 0.565 | 4.10E-07 | 0.209 | [0.128,0.289] | [NEK1] |
| rs4702 | 15 | 91426560 | A/G | 0.456 | 4.40E-07 | -0.201 | [-0.280,-0.123] | [FURIN] |
| rs374346895:AAC | 22 | 36199194 | D/I | 0.089 | 4.60E-07 | -0.372 | [-0.516,-0.227] | [RBFOX2] |
| rs1430348 | 2 | 73367058 | C/T | 0.447 | 4.70E-07 | -0.204 | [-0.283,-0.125] | RAB11FIP5--[]--NOTO |
| rs67951111 | 16 | 14493828 | C/G | 0.216 | 6.00E-07 | 0.252 | [0.153,0.351] | MKL2---[]--PARN |
| rs57771728 | 7 | 138185713 | D/I | 0.501 | 6.70E-07 | 0.201 | [0.122,0.280] | [TRIM24] |
| rs5844547 | 22 | 23460684 | D/I | 0.292 | 7.70E-07 | 0.222 | [0.134,0.311] | [RSPH14,GNAZ] |
| rs72816098 | 5 | 166098090 | A/G | 0.090 | 8.30E-07 | 0.344 | [0.207,0.481] | []---TENM2 |
| rs9922099 | 16 | 76480999 | A/G | 0.590 | 8.90E-07 | -0.203 | [-0.284,-0.122] | [CNTNAP4] |

### Supplementary Figures

#### Supplementary Figure 1. An example of what a user would see during a trial of the Vocabulary Knowledge task

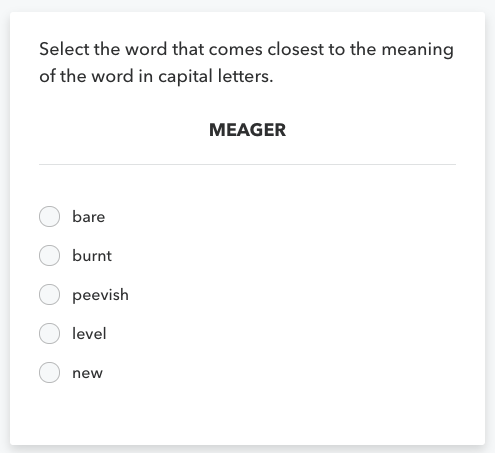

#### Supplementary Figure 2. An example of what a user would see during a trial of the Flicker Change Detection task

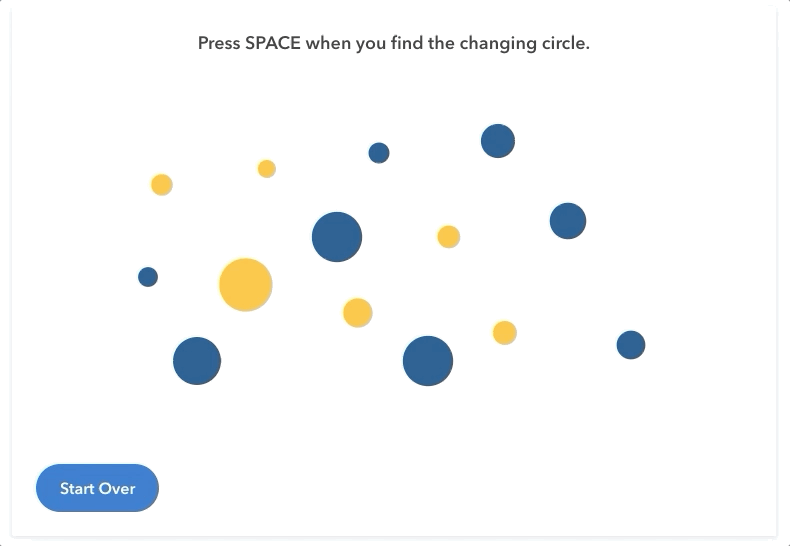

#### Supplementary Figure 3. An example of what a user would see during a trial of the Digit Symbol Substitution Test

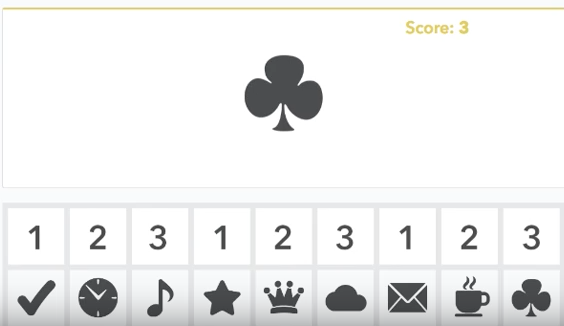

#### Supplementary Figure 4. Q-Q plot for the Vocabulary Knowledge GWAS

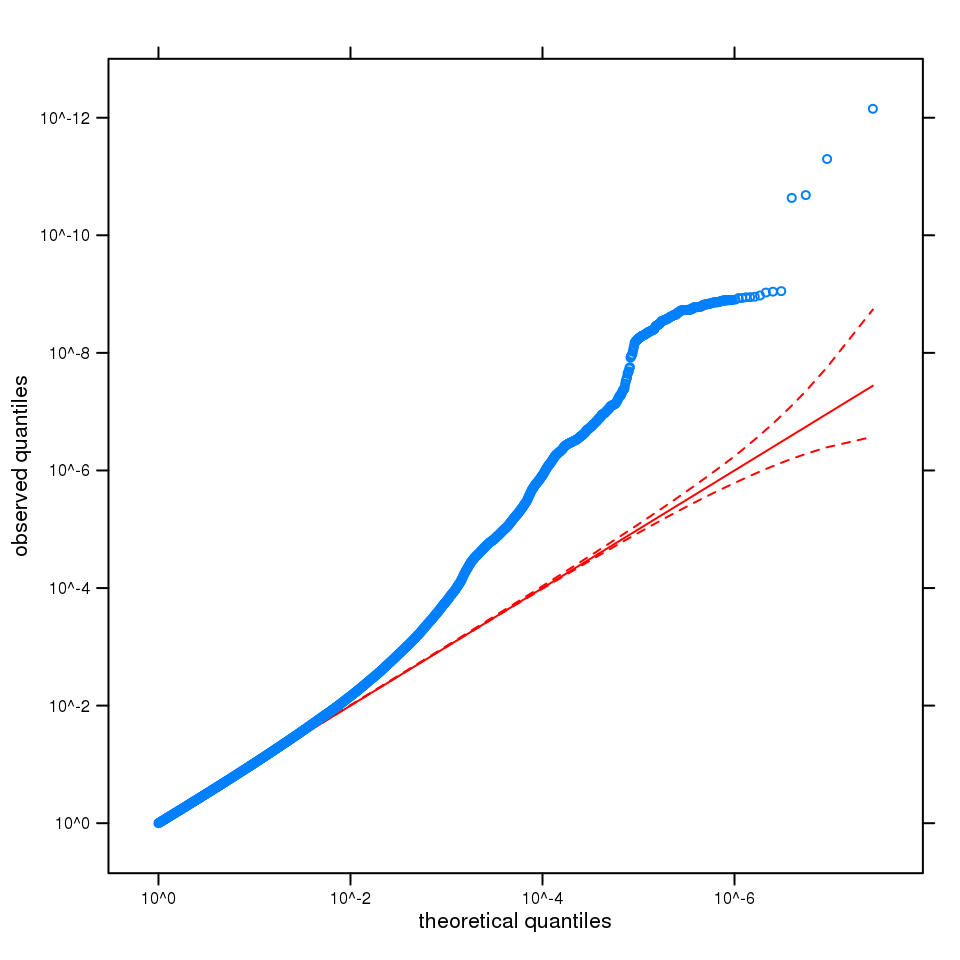

Q-Q plot of observed versus expected quantiles for the GWAS P values for Vocabulary Knowledge, where the expected distribution of P values is uniform under the null hypothesis, plotted on a log scale. A solid red line is shown with a slope of 1, and dashed red lines represent a 95% confidence envelope under the assumption that the test results are independent. The test statistics in the Q-Q plot have already been adjusted for inflation, λ=1.199. The equivalent inflation factor rescaled for a sample size of 2000 would be λ_2000_=1.002, and for 20000, λ_20000_=1.021.

#### Supplementary Figure 5. Q-Q plot for the Flicker Change Detection GWAS

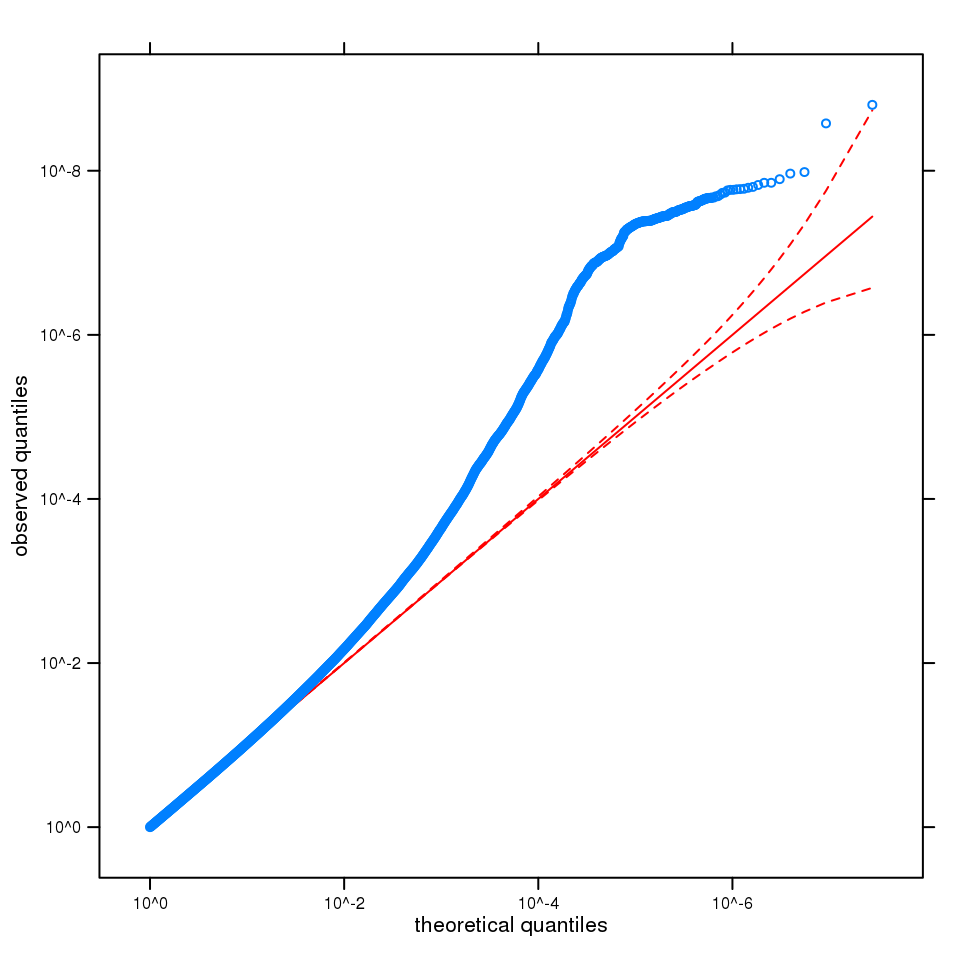

Q-Q plot of observed versus expected quantiles for the GWAS P values for Flicker Change Detection, where the expected distribution of P values is uniform under the null hypothesis, plotted on a log scale. A solid red line is shown with a slope of 1, and dashed red lines represent a 95% confidence envelope under the assumption that the test results are independent. The test statistics in the Q-Q plot have already been adjusted for inflation, λ=1.196. The equivalent inflation factor rescaled for a sample size of 2000 would be λ_2000_=1.002, and for 20000, λ_20000_=1.025.

#### Supplementary Figure 6. Q-Q plot for the Digit Symbol Substitution Test GWAS

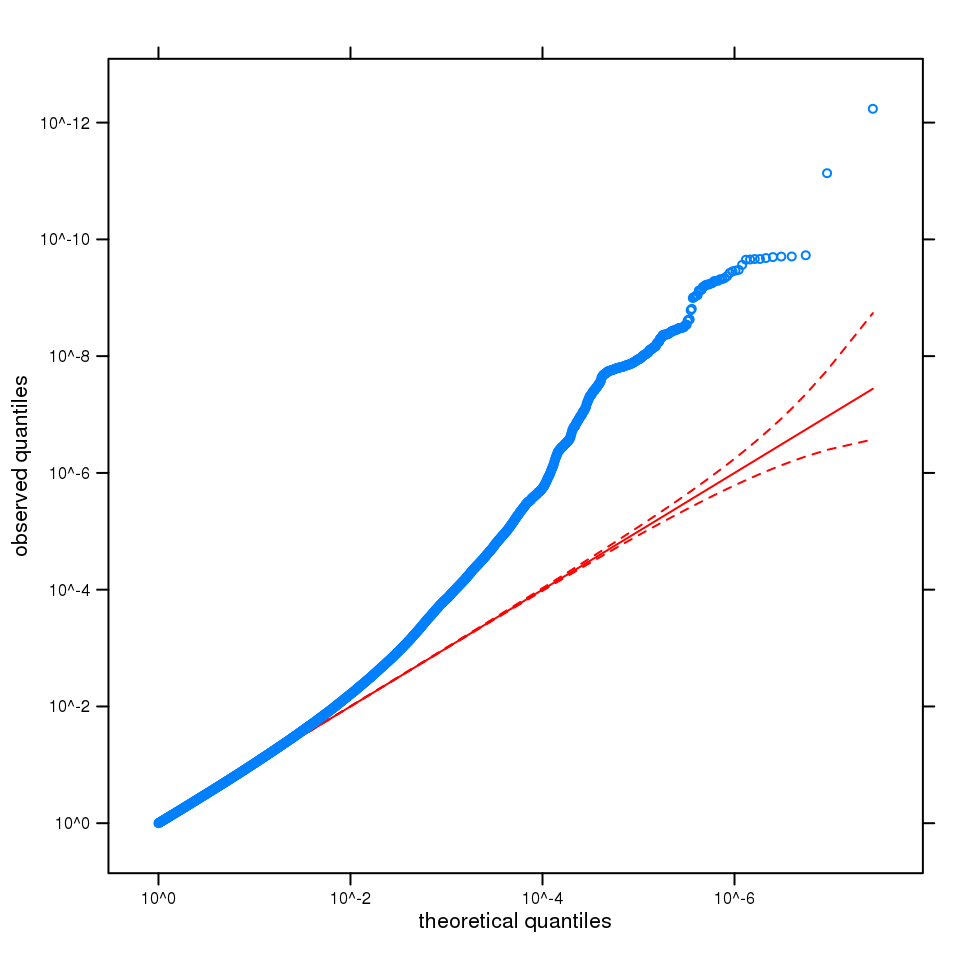

Q-Q plot of observed versus expected quantiles for the GWAS P values for Digit Symbol Substitution, where the expected distribution of P values is uniform under the null hypothesis, plotted on a log scale. A solid red line is shown with a slope of 1, and dashed red lines represent a 95% confidence envelope under the assumption that the test results are independent. The test statistics in the Q-Q plot have already been adjusted for inflation, λ=1.219. The equivalent inflation factor rescaled for a sample size of 2000 would be λ_2000_=1.003, and for 20000, λ_20000_=1.033.

#### Supplementary Figure 7. Regional association plot for rs13107325, the top hit in the Vocabulary Knowledge GWAS

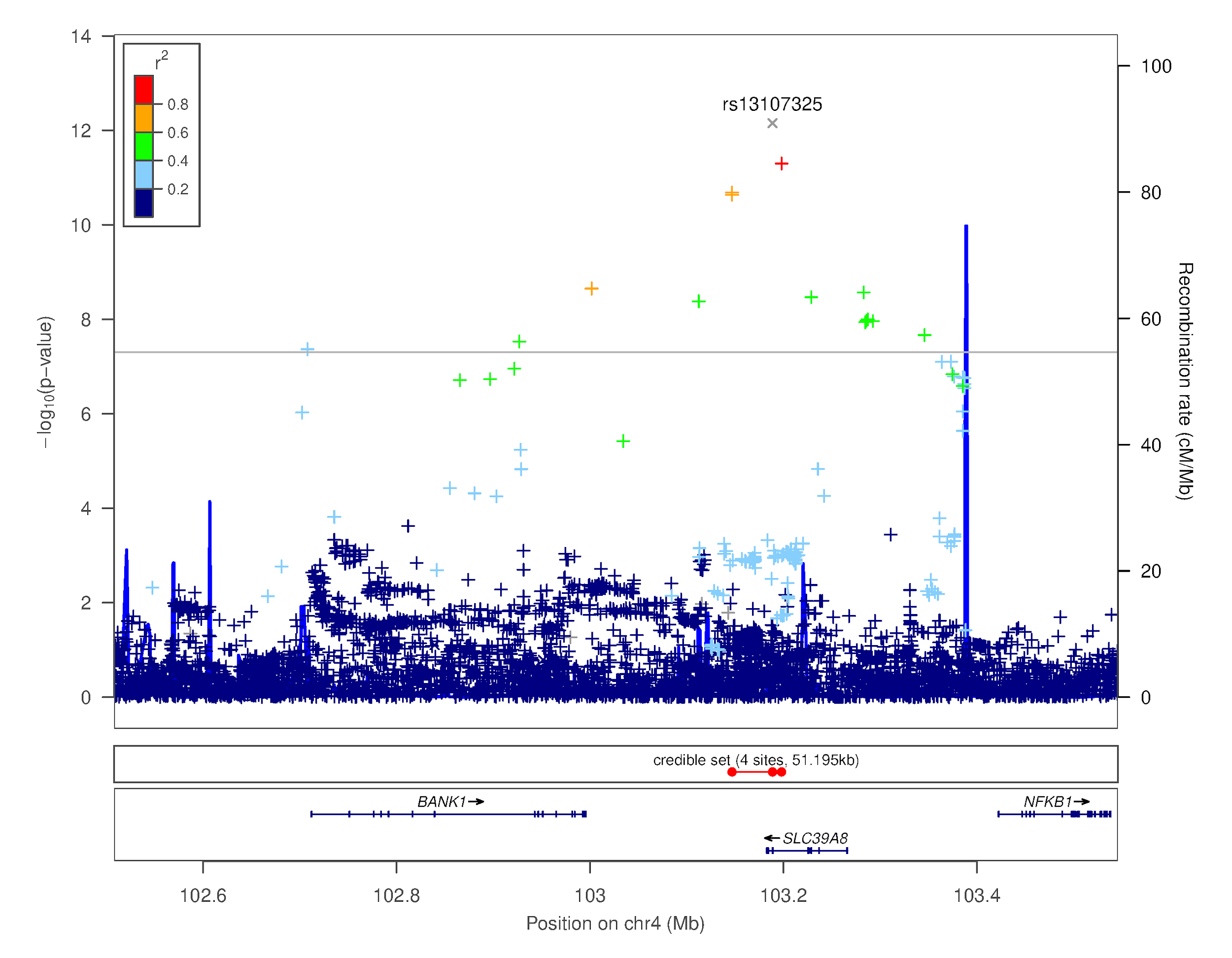

#### Supplementary Figure 8. Regional association plot for rs11316974, the top hit in the Flicker Change Detection GWAS

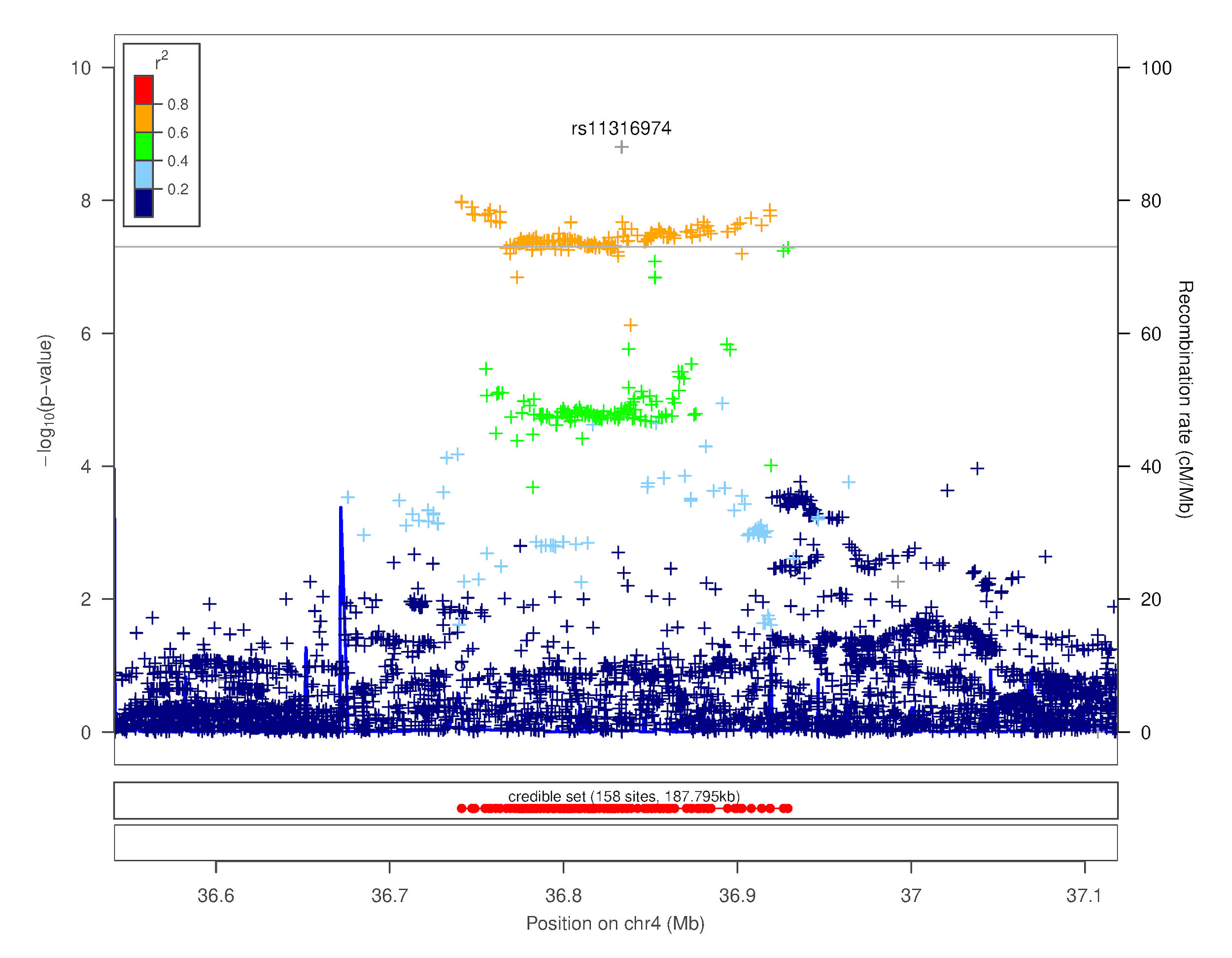

#### Supplementary Figure 9. Regional association plot for rs429358, the top hit in the Digit Symbol Substitution Test GWAS

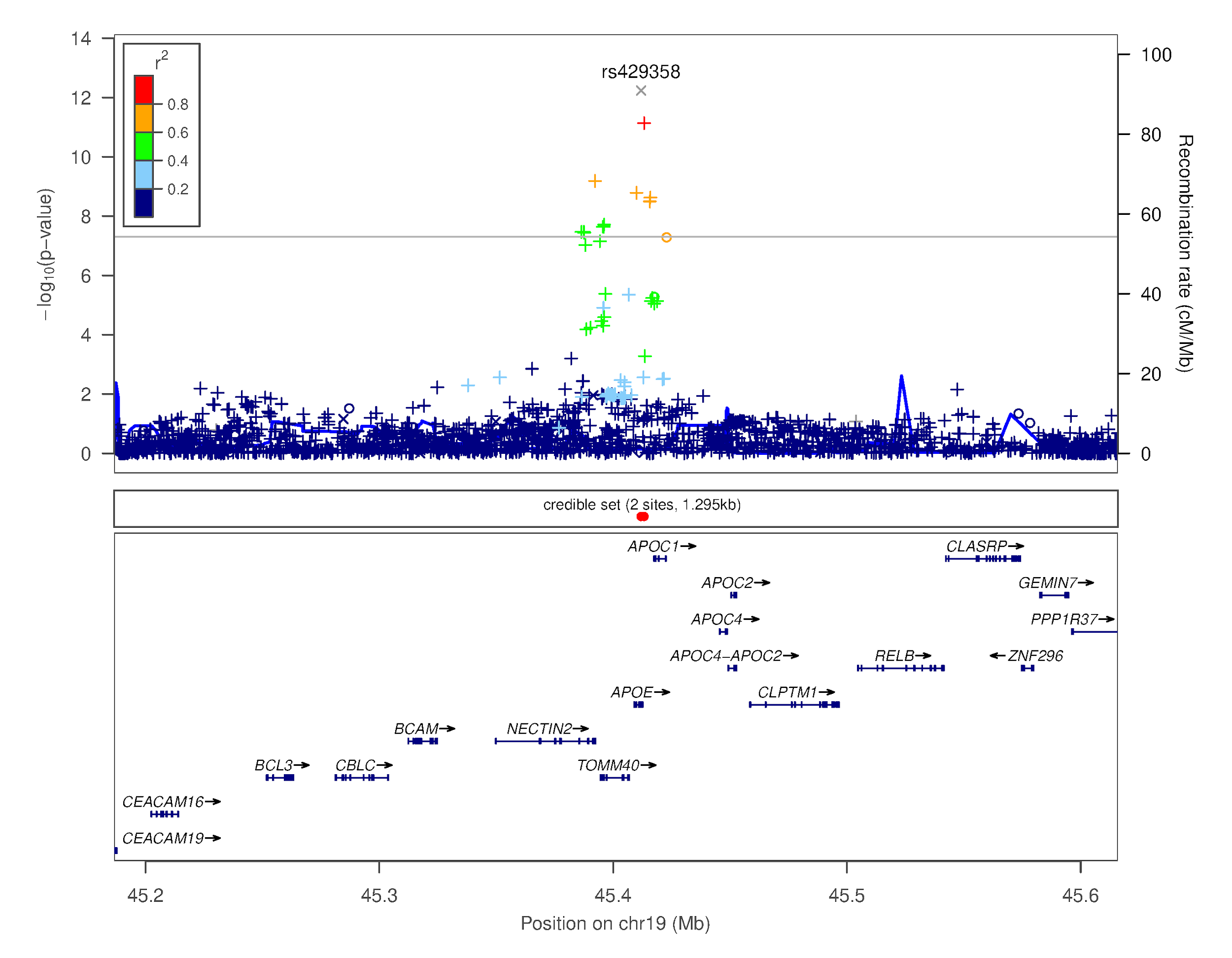
